# Genome-wide Association Studies Reveal Important Candidate Genes for the *Bacillus pumilus* TUAT-1-*Arabidopsis thaliana* Interaction

**DOI:** 10.1101/2020.05.26.117002

**Authors:** Marina Soneghett Cotta, Fernanda Plucani do Amaral, Leonardo Magalhães Cruz, Roseli Wassem, Fábio de Oliveira Pedrosa, Tadashi Yokoyama, Gary Stacey

## Abstract

The plant growth promoting bacterium (PGPB) *Bacillus pumilus* TUAT-1 is an indole acetic acid producer that can increase plant growth. Inoculation with this strain has been shown to confer greater plant tolerance to drought and saline conditions. Although the ability of TUAT-1 to enhance plant growth is well documented, little is known about what mechanisms underlie the plant response to this bacterium. Applying genome-wide association study (GWAS), we evaluated the interaction between TUAT-1 and *Arabidopsis thaliana*, screening 288 plant ecotypes for root architecture traits comparing non-inoculated and inoculated plants. Most of the ecotypes were significantly affected by TUAT-1 inoculation (66.7%) for at least one of the root traits measured. For example, some ecotypes responded positively increasing root growth while others showed reduced growth upon inoculation. A total of 96 ecotypes (33.3%) did not respond significantly to PGPB inoculation. These results are consistent with the widely reported strain-genotype specificity shown in many plant-microbe interactions. The GWAS analysis revealed significant SNPs associated to specific root traits leading to identification of several genes putatively involved in enabling the *Bacillus pumilus* TUAT-1 and *A. thaliana* association and contributing to plant growth promotion. Our results show that root architecture features are genetic separable traits associated with plant growth in association with TUAT-1. Our findings validate previous reported genes involved in *Bacillus spp*.-plant interaction, growth promotion and highlight potential genes involved in plant microbe interaction. We suggest that plant-bacterial interaction and the plant growth promotion are quantitative and multigenic traits. This knowledge expands our understanding of the functional mechanisms driving plant growth promotion by PGPB.

## 1. INTRODUCTION

Modern agricultural practice focuses intensively on crop yield through the extensive use of chemical pesticides and fertilizers, which can be detrimental to the environment. Looking to the future, agriculture needs to be more sustainable and environmentally friendly. Increasing attention is now being paid to soil microorganisms that have the ability to promote plant growth in a sustainable way. These plant growth promoting bacteria (PGPB) can colonize roots to quite high levels (≥10^8^ CFU/ gm fresh weight) without eliciting a plant defense (Faoro 2017 and Reinhold-Hurek & Hurek, 1998, 2011). PGPB commonly impact root architecture and plant health, attributing these effects to management of biotic and abiotic stresses (Vejan 2016), production of phytohormones, enhancing nutrient acquisition, and protection against pathogens and pests (Pankievicz 2015 and Pérez-Montaño 2014) Nevertheless, definitive evidence that defines the specific mechanism of PGPB-mediated plant growth promotion remains unknown (Vejan 2016).

Various plant-bacterial interactions have been intensively studied, although relatively few studies have specifically focused on determining the mechanisms by which PGPB stimulate plant growth. Genome wide association studies (GWAS), a powerful tool to analyze natural variation, have been used to study PGPB-plant interactions. For example, one study analyzed the interaction of *Pseudomonas simiae* WCS417r with *Arabidopsis thaliana* (Wintermans 2016). *A. thaliana* was one of the first non-human organisms investigated by GWAS analyses (Atwell 2010). This plant is the ideal organism for GWAS analysis, since it can be maintained as pure inbred lines through self-fertilization, which makes it possible to repeat phenotyping of genetically identical individuals (Korte and Farlow 2013).

Most PGPB belong to the Proteobacteria and Firmicutes phyla. Among firmicutes, *Bacillus spp*. is the predominant PGPB. These bacteria are spore-forming and can survive for a long time under unfavorable conditions (Radhakrishnan 2017). The strain *B. pumilus* TUAT-1, was originally isolated from rice roots, and was demonstrated to increase root growth and biomass, as well as plant nitrogen and chlorophyll content. Some *B. pumilus* can produce gibberellin and increase the nitrogen content and shoot surface area in plants (Probanza 1996). This bacterium produces significant amounts of gibberellins, more than 200 ng.ml^-1^ after 24 h (Gutierrez-Manero 2001 and Lee 2006). *B. pumilus* was also shown to promote growth and stress tolerance under conditions of drought or high salt stress and has antifungal activity (Gurav 2017 and Win 2018).

The complete genome sequence of TUAT-1 has been reported and for this strain, the production of indole acetic acid was confirmed (Okazaki 2019).

In this study, we used genome-wide association analysis (GWAS) to explore the ability of *B. pumilus* TUAT-1 to colonize and promote the growth of *Arabidopsis thaliana* seedlings. A panel comprising 288 *A. thaliana* ecotypes was used with the aim to identify traits and genomic regions associated with the plant growth promotion.

## 2. MATERIAL AND METHODS

### 1. Plant inoculation and growth promotion assay

A panel with 288 *A. thaliana* ecotypes was used to perform this study, derived from germplasm stock CS76309, maintained by The Arabidopsis Information Resource (TAIR on www.arabidopsis.org) (Berardini 2015). The plants were grown under controlled conditions at 22° C with a 16/8 h photoperiod. Arabidopsis seeds were surface sterilized and vernalized for 4 days at 4° C in the dark. Seeds were germinated in Petri dishes with ½ Murashige-Skoog (MS) medium with 0.5% sucrose (Murashige and Skoog 1962), then placed vertically in a growth chamber. After 6 days, seedlings with the same size were transferred to a new plate with ½ MS medium with 0.5 % sucrose and inoculated. For each ecotype, there were 3 plates containing 5 plants for the uninoculated controls and inoculated. The plates were placed vertically in a growth chamber for an additional 6 days. For the control plants, a drop of 25 μL of 1% saline was added to the plate approximately 3 cm distant from the root tip for each seedling. Treated plants were inoculated with a drop of 25 μL of TUAT-1 (10^8^ CFU/mL), obtained from a 10-times diluted overnight culture grown on Tryptic Soy Broth (15 g/L, TSB, Difco^™^) medium at 30°C under rotation (130 rpm).

Images were obtained 6 days after inoculation with EPSON PERFECTION V500 PHOTO scanner at 800 dpi. The root pictures were analyzed using EZ-Rhizo software (Rogers, 2018), which measured the following root traits: main root length (MRL), number of lateral roots (NLR), branched zone (BZ), total root length (TRL) and lateral root length (LRL). In total, the 5 traits were measured in 8,640 roots. Averaged root trait data was calculated with GraphPad Prism version 8.00 in Windows. Statistically significant differences between control and inoculated plants was accessed by the same program using multiple Student t test (*: p < 0.05).

The 5 trait phenotypes were measured by comparing inoculated plants to control plants. These phenotypes were calculated by Δ = A2 – A1, where A1 was the average of the control plants and A2 was the average of the inoculated plants. A phenotype heatmap was built using OLIVER (Tessmer, 2019) to compare all of the phenotypes from the 288 ecotypes and the relation between the traits.

### 2. GWAS analysis

The average (Δ) of each phenotype was used to perform GWAS analysis. Genome-wide association mapping was conducted using GWA-Portal web interface (Seren, 2018). In this study, two different data sets were used to perform GWAS analysis. The first set was composed of <u>a</u>ll <u>e</u>cotypes (ΔAE data set) that were phenotyped while the second set was composed of only those ecotypes that showed at least one trait with a significant difference between control and inoculated plants, referred here to as responsive ecotypes (ΔRE data set).

The 250K SNPs data set from GWA-Portal was used to perform the analysis. The transformations used were LOG and SQRT and the algorithm method was the accelerated mixed model. SNPs with *p values* lower than 10^-7^ were selected from the growth promotion parameters for gene identification.

### 3. Genetic trait identification

To evaluate the impact of the significant Single Nucleotide Polymorphisms (SNPs), we analyzed the location of each SNP and the allele effect on the gene using GWA-Portal (Seren, 2018). The SNPs were classified in the following categories: i) stop gained, when the base changed in the SNP introduce a stop codon inside a coding region, presumably producing a shorter protein; ii) start lost, when the base changed in the SNP occurs in a start codon, removing it; iii) non-synonymous coding, when the base changed in the SNP, also changing the codon and the coded amino acid in the protein; iv) synonymous coding, when the base changed in the SNP, changing the codon but not the coded amino acid in the protein (synonymous codon); v) untranslated region (UTR) prime, when the SNP occurs in a non-coding region, 5’ or 3’ of coding sequence into the transcribed region (mRNA); vi) intron, when the base changed in the SNP occurs in an non-coding intron sequence, and vii) intergenic, when the base changed in the SNP occurs in non-transcribed region.

Gene identification was performed with GWA-Portal and confirmed using the GBrowse tool from The Arabidopsis Information Resource (TAIR) at https://gbrowse.arabidopsis.org/cgi-bin/gb2/gbrowse/arabidopsis/. When a significant SNP (*SNP) was found inside an intergenic region (ing*SNP), linkage disequilibrium (LD) was verified in order to identify the presence of other SNPs that were possibly correlated to the ing*SNP, within 10 kb up or downstream, using the GWA-Portal (SEREN, 2018). SNPs in LD found with a correlation coefficient (r^2^) higher than 0.5 were then used for candidate gene identification.

In the next step, coding regions identified from *SNPs and LD analysis from ing*SNPs were derived for its predicted proteins and further functional analysis was performed. The Gene Ontology (GO) terms were analyzed using the GO Annotations, from TAIR (https://www.arabidopsis.org/tools/bulk/go) and gene ontology networks were built using ClueGo (Bindea 2009) to decipher functionally grouped genes.

Gene network interaction were built in order to evaluate the functional relation of the identified genes. These networks were created using identified genes, from both data sets, using Cytoscape (Shannon, 2003) with the String database version 10.5 (Szklarczyk 2019). A comparative analysis was performed between our results and *Pseudomonas simiae* WCS417r-*Arabidopsis thaliana* GWAs. (Wintermans 2016).

### 4. Candidate gene selection

Candidate genes were chosen according to the following criteria: genes identified in both data sets (AE and RE) or in both transformation analysis (LOG and SQRT), genes with allele alterations that would cause a non-synonymous coding (nsy*SNPs) or a stop codon (sp*SNP), genes with SNPs in LD with SNP 29235922 in chromosome 1, and genes from the network interaction that had a high number of interactions.

To evaluate if the candidate genes were co-regulated with other genes, we used the expression data sets in the ATTED-II (Obayashi, 2018) gene co-expression database version 9.0.

## 3. RESULTS

### 1. Response of *A. thaliana* ecotypes to TUAT-1 inoculation

In order to evaluate the growth promoting capability of *Bacillus pumilus* TUAT-1, a PGPB commercially used for rice crops in Japan, we screened 288 *Arabidopsis thaliana* ecotypes upon TUAT-1 strain inoculation. Various root traits were analyzed and then submitted to genome-wide association analysis (GWAS) to identify possible genes involved in plant-bacterial interaction.

Each of the 288 *Arabidopsis* ecotypes were evaluated for plant growth promotion by measuring root traits, comparing control and inoculated plants (supplementary table 1). TUAT-1 inoculation significantly affected root architecture traits compared to control plants, as follow: 52.7% for main root length (MRL), 14.2% for numbers of lateral root (NLR), 8.3% for branched zone (BZ), 21.2% for total root length (TRL), and 19.1% for lateral root length (LRL). Many inoculated ecotypes were significantly different from the control for more than one trait; however, only ecotype Ts-1 showed significant differences for all tested traits (Fig. 1 a). The plant response to inoculation (Sup. Fig. 1) was categorized into three groups: 1) non-responsive, 2) positive response or 3) negative response (Fig. 1e-f) according to statistical differences in trait measurements between control and inoculated plants.

**Figure 1.**
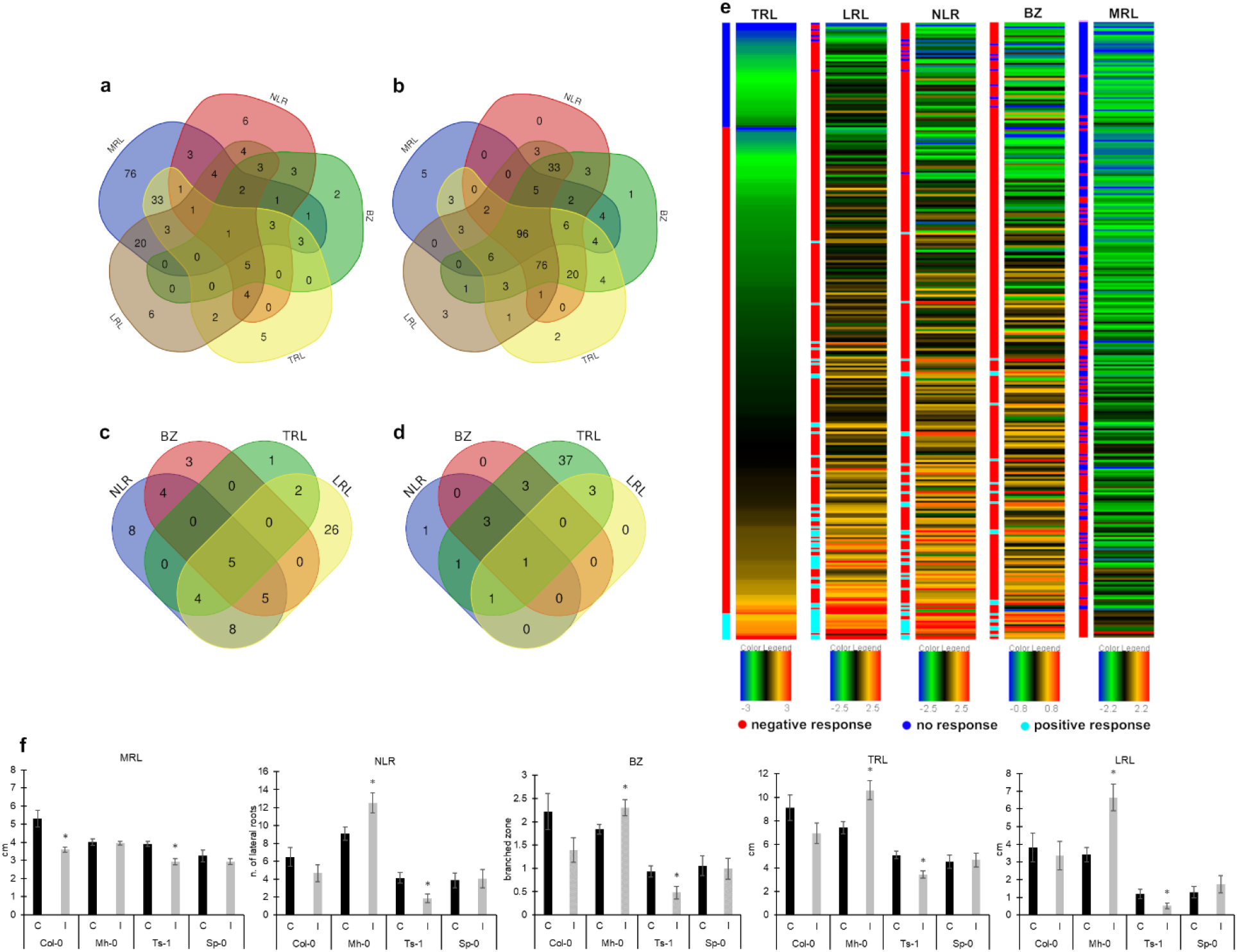
Phenotype screening from 288 A. thaliana ecotypes inoculated with TUAT-1. (a) Venn diagram of significant ecotypes response. (b) Venn diagram of no significant ecotypes response. (c) Venn diagram of positive ecotypes response. (d) Venn diagram of negative ecotypes response. (e) Heatmap of the ecotypes Δ phenotype. The heatmap was built sorting from the lowest Δ phenotype to the highest and then separated by the plant response category for TRL trait. In the following traits, the ecotypes are in the same order as the TRL trait. (f) Growth promotion analysis of the plant response types. Mh-0, Sp-0 and Ts-1 are, respectively, positive, non and negative responsive ecotypes for the traits that showed significant difference between control and inoculated plants. Col-0 is the reference ecotype.

All ecotypes inoculated with TUAT-1 showed on average a 26.2% reduction of MRL indicating that this is a general effect of this strain. The decrease of MRL was already reported in *A. thaliana* (Col-0) inoculated with *Bacillus amyloliquefaciens* UCMB5113 (Asari 2017). For the remaining traits, the tested ecotypes showed variation in plant response to inoculation (positive or negative). Positive responsive plants showed an increment of 113.2% (NLR), 264.4% (BZ), 43.3% (TRL) and 175.6% (LRL). Whilst, negative responsive plants showed a reduction of 37.8% (NLR), 40.5% (BZ), 25.6% (TRL) and 37.5% (LRL). In our study, 96 (33.3%) ecotypes showed no response to TUAT-1 inoculation for all traits evaluated (Fig. 1 b)). (Wintermans et al., 2016). Due to the observed variation in the plant response to inoculation, further analysis was applied considering two sub-sets: 1) all ecotypes (AE) and 2) only responsive ecotypes for at least one trait (RE).

The response of individual ecotypes through different traits is displayed as a heat map in Figure 1e, showing all ecotype phenotype values and their response categories (Fig. 1 e), which clearly demonstrates that most of the ecotypes analyzed showed different responses among the traits. The calculated correlation coefficients showed a positive correlation for each pair of traits in both data sets (AE and RE). The strongest correlation was found in TRL and LRL traits, with a correlation coefficient of 0.83 (AE) and 0.88 (RE). The lowest correlation coefficient was found between MRL and LRL traits, 0.13 (AE) and 0.27 (RE). We also analyzed the link between phenotype Δ values to access the phenotype correlation. Although all the trait pairs showed a positive correlation, the link between the Δ ecotype values (Supp. Fig. 2) suggested that the MRL and LRL traits, with the lowest correlation coefficient, were inversely correlated. Therefore, our data suggest that the lateral root length is the main trait that contributed the most to the measured variation in total root length.

### 2. *A. thaliana* SNPs significantly associated with *B. pumilus* TUAT-1 inoculation

Since a large number of non-responsive ecotypes were observed, in addition to analyzing that data from all 288 ecotypes (AE data set), a second data set was created, excluding the 96 (33.3%) non-responsive ecotypes, composed of the remaining 192 (67.7%) responsive ecotypes (RE data set). We decided to create the RE data set because the Δ values (differences between control and inoculated plants) were close for many non-responsive ecotypes and responsive ecotypes. However, due to the sample variation, no statistical significance was achieved in the first group.

GWAS was performed using the GWA-Portal web interface (Seren, 2018) to analyze the genetic basis of the root architecture of *A. thaliana* in response to TUAT-1 inoculation. The differences between control and inoculated plants for each evaluated trait were used as input values for AE and RE data sets. The 250K SNPs database included in GWA-Portal covered 97% and 95% of AE and RE data sets, respectively. Since the phenotype data did not fit a normal distribution (data not shown), we initially transformed the raw data using LOG transformation. The analysis was also performed from raw data transformed by SQRT transformation, resulting in a lower number of significant SNPs (data not shown). The significant SNPs found in this analysis were also identified by analysis from LOG transformation, giving more support for the selection of genes potentially involved in plant-bacterial interaction and plant growth promotion.

GWAS resulted in a list of 160 and 135 SNPs with significant p-values (<10^-7^) (Supp. Table 2) for the AE and RE data sets, respectively (Fig. 2a-b). Even with a difference of 96 ecotypes, the data sets shared 72 SNPs (Fig 2 c). Several SNPs were found to be associated with more than one trait. TRL and LRL traits shared 19 and 49 SNPs, for AE and RE data sets, respectively (Fig. 2 d and e). Even with a difference of 96 ecotypes, the data sets shared 72 SNPs.

**Figure 2.**
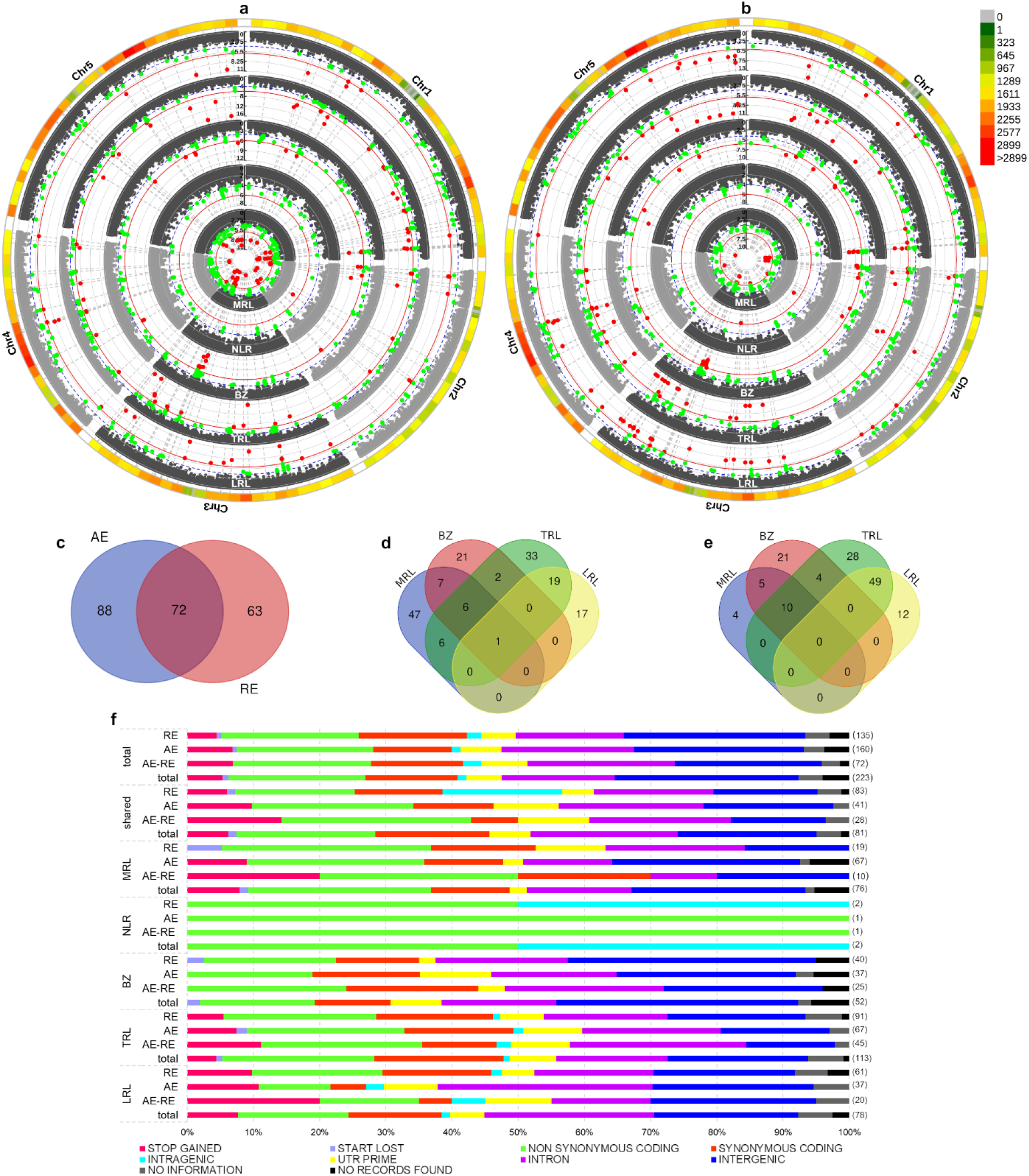
Significant SNPs identified by GWAS. (a) Circular Manhattan plot from GWAS analysis for AE data set. (b) Circular Manhattan plot from GWAS analysis for RE data set. Colorful circle around Manhattan plots shows SNPs density across the chromosomes. (c) Venn diagram of significant SNPs found in both data sets. (d) Venn diagrams of significant SNPs associated with MRL, BZ, TRL and LRL traits for AE data set. (e) Venn diagrams of significant SNPs associated with MRL, BZ, TRL and LRL traits for AE data set. NLR trait was excluded from Venn diagrams because this trait did not share SNPs with other traits. (f) SNP impact analysis. SNPs were classified as stop gained, start lost, no synonymous coding, synonymous coding, UTR prime, intron and intergenic. SNPs classified as intragenic was inside gene region, however there is no information about the allele effect. SNPs categorized as “no information” presented allele frequency, but no information about the allele effect and “not records found” SNPs had no information in the database used. The proportion of each SNP impact category was analyzed in 4 different data groups: AE data set (AE), RE data set (RE), SNPs shared by the two data sets analysis (AE-RE) and the total number of SNPs from both data sets (total). Number between parenthesis besides each bar means the number of SNPs found in that data group.

SNP location and allele effect were analyzed for all significant SNPs (*SNPs), from both data sets, to evaluate the SNP’s impact (Fig. 2 f). SNPs found in this study were mostly separated in the following categories: intergenic (ing*SNP, 27.8%), non-synonymous coding (nsy*SNP, 20.63%), intron (int*SNP, 17%), synonymous coding (sy*SNP, 13.9%), stop gained (sp*SNP, 5.4%) and UTR prime (utr*SNP, 5.4%). For those intergenic significant SNPs (ing*SNPs), the linkage disequilibrium (LD) was also analyzed, resulting in 40 and 41 SNPs associated with the ing*SNPs in AE and RE data sets, respectively. Among these new SNPs, 23 and 27 were found to be significant in the GWAS for AE and RE data sets, respectively. Including those SNPs in LD with the first set of ing*SNPs, 40% were classified as intergenic, 23,3% as intron, 13,3% as non-synonymous coding, 10% as synonymous coding, and 10% as UTR prime.

### 3. Genetic traits associated with *A. thaliana* and *B. pumilus* TUAT-1 interaction

Genes around the *SNPs classified as non-synonymous coding (nsy*SNPs), intron (int*SNPs), synonymous coding (sy*SNP), stop gained (sp*SNPs) and, UTR prime (utr*SNPs) targeted 128 genes, most predicted to result in loss of function. Linkage disequilibrium analysis of ing*SNPs led to 39 genes, from which, 24 were already identified in the GWAS analysis around significant SNPs (Fig. 3a). SNPs in linkage disequilibrium with ing*SNPs were selected according to the same classification for *SNPs found in GWAS analysis and these categories were selected because these alterations could imply possible loss of function. AE dataset analysis provided a list of 94 genes from GWAS and LD analysis and 79 genes from RE dataset. The data sets shared 47 genes (Fig. 3 b) and, in total, the final gene list was composed of 143 genes (Supp Table 2).

**Figure 3.**
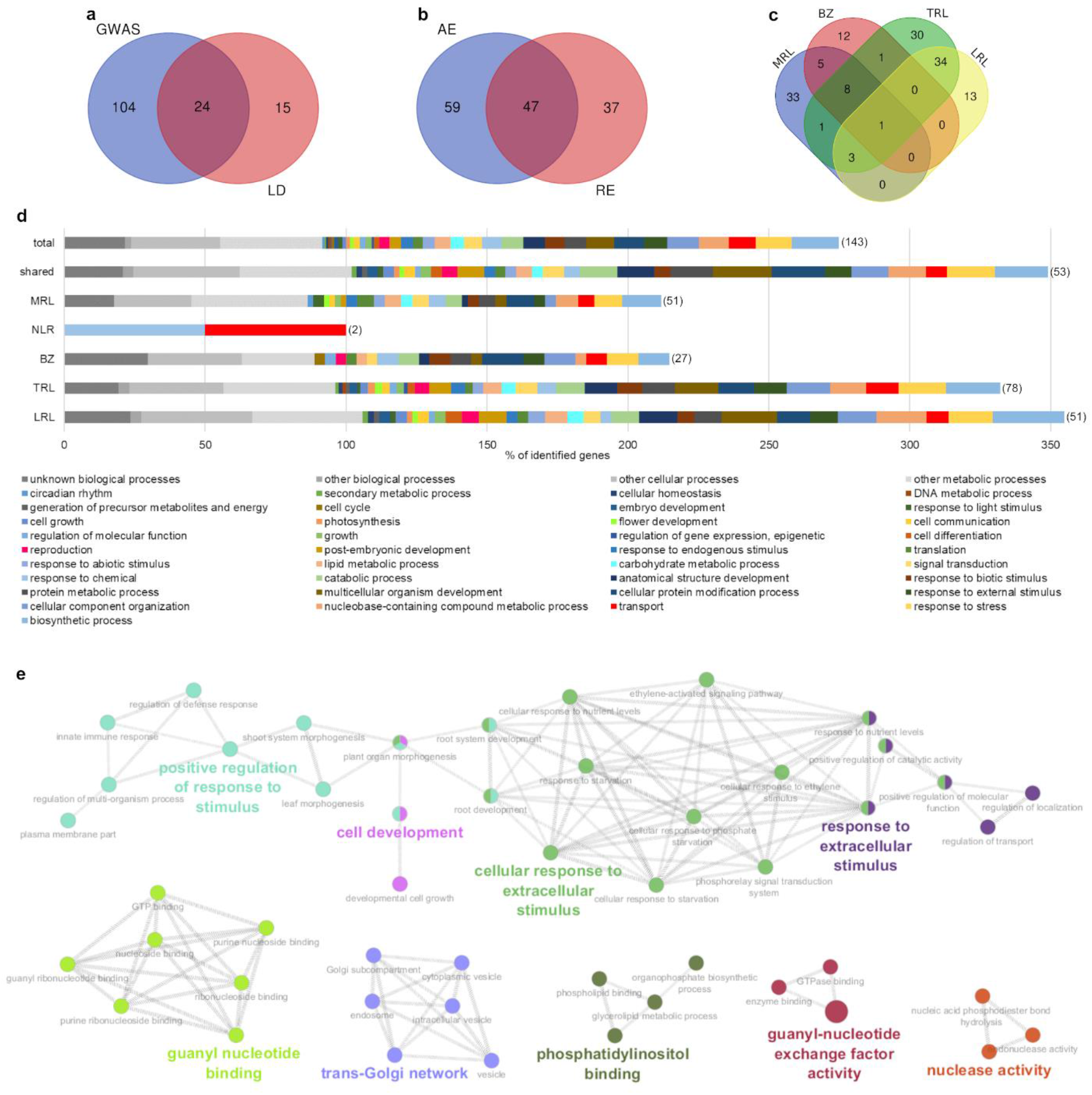
Genetic traits identified by GWAS. (a) Venn diagram of identified genes from GWAS and LD analysis in both data sets. (b) Venn diagrams of identified genes in AE and RE data set GWAS and LD analysis. (c) Venn diagrams of identified genes associated with MRL, BZ, TRL and LRL traits for identified genes in both analyses. NLR trait was excluded from Venn diagrams because this trait did not share genes with other traits. (d) GO terms for biological processes for each trait, total number of identified genes in all the traits (total) and genes associated with more than one trait (shared). Number between parenthesis represent the number of genes in that data group. The sum of the percentage is higher than 100% because the genes can belong to more than one biological process at the same time. (f) GO network built with all identified genes for biological processes, molecular function and cellular component. Colorful names are the group leading terms.

Among all identified genes, 37.1% were associated with more than one trait. TRL and LRL were the traits that shared the most genes, 34. These results are consistent with the phenotype correlation between the traits, where TRL and LRL were the traits with the strongest positive correlation. Even though many genes were associated with more than one trait, most of the genes were associated with only one trait (62.9%). Only one gene was associated with 4 traits (*MOD1*), which encodes an enoyl-acyl carrier protein (ACP) reductase. Therefore, in this study, most of the genetic traits are associated with specific genetic regions that contribute to whole root architecture in general and genetic separable traits associated with the plant growth.

In order to have a general view of which physiological processes that appeared to be impacted by inoculation, we evaluated the GO terms for biological processes (Fig. 3 d) of the encoded proteins identified. Most of the predicted functions of the various identified genes were classified as “other or unknown cellular processes”, “other metabolic or cellular processes”, or “biosynthetic processes”. Among the identified biological processes, for MRL, a higher percentage of genes belonged to stress response (9.8%) and cellular protein modification processes (9.8%). For NLR we identified only 2 genes, the first one was involved in transport and the second with chemical response. For the traits BZ, TRL and MRL, genes were mostly classified as cellular protein modification processes (14.8%), response to stress (16.7%) and, multicellular organism development (19.6%). Genes associated with more than one trait were mostly involved in multicellular organism development (20.7%), while those genes related to all traits were mostly categorized in the stress response group (12.6%). This analysis showed that biological processes were specific to particular traits instead of processes affecting all traits.

GO terms were also analyzed for the total number of genes trough ClueGo plug-in (Bindea 2009). Genes were analyzed for biological processes, molecular function and cellular components and grouped according to leading terms in a network. The biggest network was composed of 26 genes and 4 leading terms: positive regulation of response to stimulus, cell development, cellular response to extracellular stimulus and response to extracellular stimulus. Smaller networks were also provided by this analysis and the leading terms were: guanyl nucleotide binding, trans-Golgi, phosphatidylinositol binding, guanyl-nucleotide exchange factor activity and nuclease activity (Fig. 3 e).

To investigate whether the genes identified in this study had any possible functional connection, we constructed gene interaction networks (Sup. Fig. 3 a). Edges in the networks represent different types of interactions: known interactions from curated databases or experimentally determined interaction, predicted interactions based on gene neighborhood, gene fusions or co-occurrence, genes that were reported in the same study, co-expressed genes and protein homology. The networks were built with all identified genes from the AE and RE data sets. From a total of 143 identified genes, 37 were connected in some way in 10 networks, 6 of them including only interaction of a pair of proteins.

From the 37 connected genes, 20 were co-expressed, forming one group composed of 10 genes, and 5 pairs of co-expressed genes. Group 1 was composed of AT3G28500, AT2G18110, AT1G77940, AT1G77750, *EIF3E, MTSSB*, AT3G11964, AT3G16840, AT1G77800 and *PSD*. This group appears to be related to protein metabolism in general and involved in important plant traits as cell proliferation and elongation, female gametophyte and plant development, shoot meristem apical activity and leaf initiation (Sup. Fig. 3 a and Supp. Table 3). Among the 5 pairs of co-expressed genes, 3 showed interesting activities as response to phosphate deficiency (pair 3), immune response (pair 4) and transport (pair 5).

### 4. Selection of candidate genes responsible for plant growth promotion

Although the results revealed several interesting genes associated to plant phenotypic traits, we applied a more restrictive criteria in order to select a few genes more likely involved in plant response to inoculation and growth promotion. First, we selected genes identified in both data sets (AE and RE), then we looked for genes with significant SNPs (*SNPs) in both data transformation analyses (LOG and SQRT); next, we selected genes with allele alterations that would cause a non-synonymous coding or a stop codon change. We also searched for genes with *SNPs in linkage disequilibrium with SNP 29,235,922 on chromosome 1, since this SNP was in a region with a high density of significant SNPs. Finally, we chose a hub gene from the interaction networks, since a high number of interactions indicate a gene potentially affecting many biological processes. These criteria produced a list of 11 candidate genes highly likely to be related to plant-bacterial interaction and plant-growth promotion (Table 1).

**Table 1.** Candidate genes around the significant SNPs. For each gene, the p-values of the SNPs are shown. The possible alteration function of the allele is represented in the table. The last column shows the function of the gene. *Chromosome **Data set ***Transformation type of the raw data.

| Gene Identifier | Gene Name | Position | Chr.* | Dat.** | Transf.*** | p-value |  |  |  | Alteration Function | Gene Function |
| --- | --- | --- | --- | --- | --- | --- | --- | --- | --- | --- | --- |
|  |  |  |  |  |  | MRL | BZ | TRL | LRL |  |  |
| AT4G17730 | SYP23 | 9865729 | 4 | AE | LOG |  |  | 7.57E-08 | 4.15E-10 | missense | transport |
|  |  |  |  | RE | LOG |  |  | 6.10E-07 | 1.41E-09 |  |  |
|  |  |  |  | AE | SQRT |  |  |  | 1.53E-07 |  |  |
|  |  |  |  | RE | SQRT |  |  |  | 1.72E-07 |  |  |
| AT5G44210 | ERF9 | 17807267 | 5 | RE | LOG |  |  | 3.91E-11 | 4.74E-09 | missense | ethylene response factor |
| AT2G21170 | TIM | 9072376 | 2 | RE | LOG |  |  | 2.35E-10 | 9.71E-13 | silent | plastid isoform of triose phosphatase isomerase |
|  |  |  |  | RE | SQRT |  |  |  | 1.32E-07 |  |  |
| AT5G64510 | TIN1 | 25786075 | 5 | AE | LOG |  |  | 1.44E-16 | 4.74E-11 | nonsense | plant specific endoplasmic reticulum stress inducible gene |
|  |  |  |  | RE | LOG |  |  | 3.91E-11 | 4.74E-09 |  |  |
| AT3G58340 |  | 21590066 | 3 | AE | LOG |  |  | 1.44E-16 | 4.74E-11 | nonsense | abiotic stress response |
|  |  |  |  | RE | LOG |  |  | 3.91E-11 | 4.74E-09 |  |  |
| AT3G61350 | SKIP4 | 22704328 | 3 | AE | LOG |  |  | 1.44E-16 | 4.74E-11 | nonsense | involved in the regulation of circadian clock |
|  |  |  |  | RE | LOG |  |  | 3.91E-11 | 4.74E-09 |  |  |
| AT1G77760 | NR1 | 29235922 | 1 | AE | LOG | 4.47E-11 |  |  |  |  | nitrate assimilation |
|  |  |  |  | RE | LOG | 2.16E-07 |  |  |  |  |  |
|  |  |  |  | AE | SQRT | 9.01E-08 |  |  |  |  |  |
| AT1G77800 | PRE-2 | 29254754 | 1 | AE | LOG | 7.40E-09 |  |  |  | missense | regulation of cell elongation and plant development |
|  |  | 29255670 |  |  | LOG | 2.24E-07 |  |  |  | missense |  |
| AT1G77860 | KOM | 29284188 | 1 | RE | LOG | 1.07E-07 | 6.90E-10 | 4.45E-07 |  | missense | rhomboid-related intramembrane serine protease |

Candidate genes were found nearby *SNPs in all traits, except for NLR, and were distributed across all 5 *Arabidopsis thaliana* chromosomes. A subset of 6 out of 11 candidate genes (*SYP23, ERF9, PRE2, KOM* and *ATG14A*) presented SNPs in which the allele alteration would cause a non-synonymous coding change, leading to a missense mutation. The *TIN1*, AT3G58340, and *SKIP4* genes are located flanking SNPs in which the allele alteration would insert a stop codon.

The *PRE-2, KOM, ATG14A*, and *DJC65* genes were found flanking SNPs in linkage disequilibrium with SNP 29,235,922 on chromosome 1 (also identified as significant in the GWAS). This SNP is located in one out of three prime untranslated regions inside the candidate gene NR1. The *DJC65* and *TIM* genes were flanking a SNP in which the allele alteration would cause a synonymous coding, and hence, a silent mutation. However, these genes were selected as candidates since the *DJC65* gene matched the criteria described above and the *TIM* gene is a hub in the gene interaction network connected to several other genes (Supp. Fig. 3).

Next, for the selected candidate genes, the possible influence of allele alteration caused by the SNPs on the phenotype (as calculated by the difference of inoculated and non-inoculated plants for each trait), was evaluated using the SNP Viewer tool from GWA-Portal (Seren, 2018). The allele effect was evaluated in both data sets, AE and RE, but only in the RE data set the allele alteration showed an effect on the phenotype (Supp. Fig. 5 and 6).

For candidate genes functionality, we verified its possible co-regulation with other genes throughout the analysis of co-expression using experimental data sets available in ATTED-II (Obayashi 2018). We analyzed the 3 most co-expressed genes with each candidate gene. Co-expression was observed for all, except for the AT3G58340 candidate gene (Supp. Table 4). The allele effect analysis showed allele alteration in AT3G58340 (SNP 21,590,066 located at chromosome 3), had an effect on the TRL and LRL traits. This belongs to a TRAF-like family protein, tumor necrosis receptor-associated factor. TRAF proteins are molecular adaptors that regulate innate and adaptive immunity, development, and abiotic stress responses (Huang 2016).

The allele effect analysis showed that allele alteration in gene *SYP23* through SNP 9,865,729 on chromosome 4 influenced the LRL trait (Supp. Fig. 6). *SYP23* codes for a protein belonging to a subfamily of Qa-SNAREs (soluble N-ethylmaleimide-sensitive factor attachment protein receptors), responsible for protein trafficking between pre-vacuolar compartments and vacuoles. It includes a SNARE motif, required for interaction with transmembrane domain anchor SNARE proteins at vesicular membranes or target organellar membranes (JAHN and SCHELLER, 2006). *SYP23* is co-expressed with genes: i) *CKA1*, an α subunit of CK2 that regulates the circadian-clock associated 1 (*CCA1*) gene (SUGANO, 1998), involved in circadian rhythm and ribosome biogenesis in eukaryotes (KEGG pathways); ii) *WAV2*, that negatively regulates stimulus-induced root bending through inhibition of root tip rotation (Mochizuki 2005) and; iii) *VAMP724*, a gene that forms SNARE complexes with *SYP123* and *SYP132* for root hair elongation (Ichikawa 2014).

The allele alteration in *ERF9* gene through SNP 17,807,267 at chromosome 5 affected TRL and LRL traits (Supp. Fig. 6). The gene is an ethylene response factor (ERF), integrating several hormonal pathways and directly responsible for the transcriptional regulation of several jasmonate (JA)/ethylene (ET)-responsive defense genes. ERFs represent the last layer of regulation in the expression of JA/ET-responsive defensive genes (Huang 2016) and act as negative regulator of plant defense mechanisms (Maruyama 2013). *ERF9* is co-expressed with a gene coding a protein containing a domain of unknown function (*DUF241*), Auxin-Regulated Gene Involved in Organ Size (*ARGOS*) and *HIPP26* genes. The *DUF241* is a conserved domain found in proteins from many different groups of plants (*Arabidopsis*, *Oryza*, *Triticum*, *Glycine*, *Gossypium*, and many more), but with unknown function. The *ARGOS* gene expression is highly induced by auxin and is involved in the regulation of cell proliferation during organ development, hence this gene plays an important role in plant growth and development (Hu 2003). The *HIPP26* gene encodes for a heavy metal binding protein, and its overexpression in *A. thaliana* enhances toleration to Cd^33^, suggesting its role in detoxification of this metal in plants (Tehseen 2010).

The allele alteration in triosephosphate isomerase (*TIM*) coding gene by variation in SNP 9,072,376 on chromosome 2 affected the LRL trait (Supp. Fig. 6). The gene encodes for a plastid isoform of triosephosphate isomerase and plays a critical role in the transition from heterotrophic to autotrophic grow in plants. Mutation in this gene show accumulation of dihydroxyacetone phosphate and methylglyoxal, the latter probably responsible for delaying transition to autotrophic growth (Chen & Thelen 2010). The *TIM* gene is also co-expressed with *VTE3*, which encodes proteins involved in the methylation step of plastoquinone biosynthesis and vitamin E biosynthesis (Z. Cheng, 2003); *PGK1*, a phosphoglycerate kinase localized exclusively in the chloroplasts of photosynthetic tissues and co-regulated with other phosphoglycerate kinases to optimize plant growth (Rosa-Téllez 2018); and *RLP4*, a cell surface receptor involved in plant development (Wang 2008).

Allele effect analysis showed that there was an allele alteration in SNP 25,786,075 on chromosome 5 within the gene *TIN1* (tunicamycin induced 1 – a plant specific endoplasmic reticulum stress inducible gene) that affected the TRL and LRL traits (Supp. Fig. 6). Mutation in this gene affects the secretion of proteins and lipids, culminating in a visibly altered pollen surface (Iwata 2012). *TIN1* is co-expressed with *TMS1* (thermosensitive male sterile 1), a gene involved in thermotolerance in pollen tubes and response to heat shock treatment in seedlings (Ma 2015); *BF1C*, a regulator of thermotolerance, which encodes a protein that accumulates rapidly and is localized in the nuclei during heat stress (Suzuki 2008); and *DNAJ*, which encodes a chaperone belonging to heat shock protein family involved in response to various environmental stresses.

The allele alteration in *SKIP4* gene through SNP 22,704,328 on chromosome 3 affected the TRL and LRL traits (Supp. Fig. 6). This gene encodes a protein that interacts with *SKIP1*, a splicing factor involved in the regulation of circadian clock (WANG, 2012), and is co-expressed with AT5G44080, *NF-YB13* and AT5G37930 genes. AT5G44080 encoded protein is involved in the plant hormonal signal transduction (KEGG pathways) and belongs to the basic-leucine zipper (bZIP) transcriptional family that regulates diverse biological processes, such as pathogen defense, light and stress signaling, seed maturation, and flower development (Jakoby 2002). In addition, *NF-YB13* encodes a transcriptional factor, and its sub-units are involved in photoperiod-regulated flowering (Siefers 2009). Finally, AT5G37930 encoded protein also belongs to the TRAF-like protein family, as the candidate gene AT3G58340.

NR1 gene, identified through SNP 29,235,922 on chromosome 1, encodes the cytosolic minor isoform of nitrate reductase (NR), playing a role in the first step of nitrate assimilation, contributing about 15% of the nitrate reductase activity in shoots. This gene is co-expressed with *NR2, NIR1* and AT5G26200, with the former also encoding a nitrate reductase, while the second one encodes a nitrite reductase (NIR), involved in the second step of nitrate assimilation, and the last gene belongs to a mitochondrial substrate carrier family.

*PRE-2* is one of six paclobutrazol-resistance genes in *A. thaliana*, encoding proteins that act in the growth-promoting transcriptional network, coordinating the growth of floral organs and contributing to successful autogamous reproduction (Shin 2019), and possibly also with a regulatory role in gibberellin-dependent development in *A. thaliana (Lee 2006). PRE-2* is co-expressed with *PHYE*, encoding a phytochrome E acting in the regulation of shade avoidance (Devlin 1998). *TAF1*, a transcription initiation factor that works in plant resistance to genotoxic stress and also involved in the DNA damage response (Waterworth 2015), and *SUS1*, a sucrose synthase.

*KOM* is a RHOMBOID-like protein located in the envelope of chloroplasts and chlorophyll-free plastids. Plants mutated in this gene show a reduction of fertility and aberrant floral morphology (Thompson 2012). This gene is co-expressed with genes AT2G25565, AT5G08090 and AT3G57440. AT2G25565 is a C3HC4-type RING finger protein that is well studied in *Arabidopsis*. Various RING finger proteins are involved in photomorphogenesis, light signaling, secretory pathway, peroxisome biogenesis, N-end rule pathway, chromatin modifications and stress tolerance (Ma 2009). Much less studies are the AT5G08090 and AT3G57440 genes, which are described as encoding a transmembrane and hypothetical protein, respectively.

ATG14 gene encodes for an autophagy-related protein that works on the induction of autophagy and autophagosomes formation, being co-expressed with genes *G18F*, an *A. thaliana* homolog of yeast ATG18 (protein required for autophagosome formation in Arabidopsis (Xiong 2005), *RGLG2*, a gene encoding a RING domain ubiquitin E3 ligase that negatively regulates the drought stress response through the ethylene response factor 53 (ERF53; Cheng et al. 2012), and *S6K2*, a S6 kinase, which is a conserved component of signaling pathways that control growth in eukaryotes. *S6K* negatively regulates cell division and can influence the rate of polyploidization and lead to chromosome instability (Henriques 2010).

*DJC65* encodes a J domain-containing co-chaperone that recruits heat shock Hsp70 chaperones. The gene is down-regulated by cold stress (Chiu 2013), and co-expressed with genes *STN7*, *SRX* and *HAD*. *STN7* is a thylakoid protein kinase involved in the control of energy allocation in the photosystems in response to light quality and is required for phosphorylation and migration of light harvesting complex II proteins and long-term alterations in thylakoid composition (Pesaresi 2009). *SRX* encodes a sulfiredoxin that participates in the signaling mechanism in response to photooxidative stress (Rey 2007). *HAD* belongs to the haloacid dehalogenase-like hydrolase (*HAD*) superfamily, which includes a diverse substrate specificity and is involved in the enzymatic cleavage by nucleophilic substitution of carbon–halogen bonds and hydrolytic enzyme activities, including phosphatase, phosphonatase and phosphoglucomutase reactions (Koonin & Tatusov, 1994).

As detailed above, the results of the GWAS analysis predict a variety of potential candidate genes that can differentially affect *A. thaliana* root traits and overall plant physiological processes. Hence, impacting the function of these genes by bacterial inoculation likely explains the phenotypic changes detected in the various ecotypes.

## 4. DISCUSSION

The interaction between plants and bacteria can be highly beneficial for both organisms. Different taxonomic groups of bacteria have been reported to promote the growth of many different plants, including agricultural crops. Studies also demonstrated specificity among plant-bacterial interactions, showing host selectivity for specific bacterial strains or microbial communities. For example, analysis of 27 maize inbred lines across different fields, showed that a small but significant fraction of variation in microbial diversity could be attributed to host genetics (PEIFFER, 2013). *Bacillus pumilus* strain TUAT-1 was isolated in Japan and demonstrated as a plant-growth promoting bacteria (PGPB) for rice, recently released as a commercial inoculant for this crop (Win 2018). TUAT-1 can increase root and shoot fresh weight, number of crown roots, number of lateral roots and total lateral root length in rice (Ngo 2019). In our study, we evaluated the response of 288 *A. thaliana* ecotypes to TUAT-1 inoculation and demonstrated, for the first time, that TUAT-1 is able to promote the growth of *A. thaliana* in a strain-genotype specific way. Given the wide range of resources available for research with Arabidopsis, our results are clearly relevant for future investigations of this plant-bacterial interaction, especially if we want to better understand the molecular mechanisms involved. Five root architecture traits were analyzed, including main root length (MRL), number of lateral roots (NLR), branched zone (BZ), total root length (TRL) and lateral root length (LRL). From the results for plant inoculation tests, the plant response was classified in three categories: positive, negative and no response. In contrast, a study conducted with the *A. thaliana* inoculated with *Pseudomonas simiae* WCS417r showed responses for all tested accessions. In this case, all ecotypes responded positively for shoot fresh weight and number of lateral roots. In the same study, the authors classified the responses in main root length ranging from significant decreases to significant increases (Wintermans 2016).

Using the GWAS approach, we were able to identify several associations between plant genotype vs. plant phenotype, possibly involved in the plant response to inoculation and growth promotion. A total of 143 genes were identified to one or more (37.1%) of evaluated plant traits. Specially, TRL and LRL traits showed the highest number of associated genes in common (34). Particularly, MOD1 was the only gene around a significant SNP associated with 4 different traits. Other GWAS analysis of plant-PGPB interaction, were not able to identify genes associated to more than one trait (Kamfwa 2015 and Wintermans 2016). MOD1 encodes an enoyl-acyl carrier protein (ACP) reductase, which is a subunit of the fatty acid synthase complex that catalyzes the *de novo* synthesis of fatty acids. Mutants in this gene have decreased enzymatic activity, impaired fatty acid biosynthesis and a decreased amount of total lipids, which leads to effects on plant growth and development and causes premature death (Mou 2000). The association of *Bacillus* with lipid biosynthesis was already reported, showing that treatment with *B. subtilis* GB03 increased lipid synthesis and alleviate lipid peroxidation and oxidative stress under high salt condition (Han 2014). Corwin (2018) studied the association between the plant pathogenic fungus, *Botrytis cinereal*, and *Arabidopsis* and observed 31% of the genes identified by GWAS were associated with two lesion traits. However, most of the root architecture characteristics were genetically separable traits associated with plant growth (Corwin 2018). These results are consistent with our GO terms analysis, where we found biological processes specific to each measured phenotype and not global networks that impacted all phenotypes.

Among the identified genes, there are important genes related to the alleviation of abiotic stress. For instance, the pair of co-regulated genes *RDUF2* and *BCS1*, which are related to drought and oxidative stress (Ho 2008 and Kim 2012). In addition, the *MOD1* gene can be related to abiotic stress attenuation, since it catalyzes fatty acid syntheses. The increase in lipid syntheses to avoid lipid peroxidation is one of *Bacillus* mechanisms for oxidative stress alleviation (Han 2014). Genes that are co-expressed with the candidate genes are also involved in abiotic stress response, such as HIPP26 gene, involved in Cd detoxification in plants, while the *SRX* gene participates in the signaling mechanism in response to photooxidative stress (Rey 2007 and Tehseen 2010). Furthermore, there are several genes identified in this study, or co-expressed with them, that are involved in thermotolerance, such as *DJC65, TMS1, DNAJ*, and *TMS1* (Chiu 2013, Ma 2015 and Suzuki 2008). In our study, we also found a potential PGPB interaction gene, *ERF9*, which is an ethylene response factor that negatively regulates plant defense (Huang 2016). In addition, the pair of co-regulated genes *ATMIN7* and *GSL05* are also related with the plant immune response (Deslandes 2012 and Sanmartín 2020). Another important finding is the pair of co-expressed genes AL6 and UBP14, which are involved in the phosphate starvation response (Chandrika 2013 and Li 2010). The importance of this is that previous reports showed that bacterial synthetic communities can simultaneously enhance the phosphate starvation response (PSR), as well as plant nutritional stress response, immune system function and maintenance of microbial assembly (Castrillo 2017). Furthermore, there is a link between host responses and fungal colonization and phosphate availability, where it was shown that the phosphate availability impacts the host response (Hacquard 2016).

We also compared our results with the previously published data characterizing the *A. thaliana* response to inoculation with *Pseudomonas simiae* WCS417r (Wintermans 2016) and created a gene interaction network for genes found in both studies (Sup. Fig. 3). Although the genes identified in both studies are not identical, there are clear connections within the gene interaction network, suggesting they are linked by a common biological process. A sub-set of 10, out of 37 genes identified for WCS417r were connected in some way with genes from TUAT-1 and *A. thaliana* associated genes and 9 of WCS417r associated genes are co-expressed with TUAT-1 associated genes. This comparison suggests that, although plant-bacterial interaction and growth promotion does not appear to be regulated by the same genes in different associations, similar plant physiological processes may be impacted regardless of which PGPB is involved. In agreement, Wintermans et al. (2016) found no common candidate genes for the PGPB-mediated increase in shoot fresh weight and changes in root architecture for *A. thaliana* inoculated with *Pseudomonas simiae* WCS417r, but found genes related by biological process.

In conclusion, this study showed that *Bacillus pumilus* TUAT-1, a rice-growth promoting bacteria, already used as a commercial inoculant in Japan, is also able to promote growth in *Arabidopsis thaliana* in a strain-genotype specific manner. This is an important finding since *A. thaliana*, a model organism with a large amount of resources available, can now be exploited for more detailed studies of the molecular mechanisms that underlie plant growth promotion. The results clearly identify candidate genes for further validation, related to plant defense and abiotic stress alleviation. These genes are likely to be associated to the plant response to bacterial inoculation, also linked to genes identified in other studies (Wintermans 2016). Our results corroborate the *Bacillus spp*. growth promotion capability (Asari 2017; Win 2018; Win 2020), which is broadly reported. Here we present further insights to the molecular mechanism involved in PGPB. Finally, although it is tempting to always look for simple explanations to explain PGPB plant growth promotion, our results argue that, similar to other plant-microbe interactions, complex and interactive mechanisms are involved. Our study suggests that plant growth promotion by bacteria is likely a quantitative, multigene trait that can be better understood for future challenges for breeding crop plants that can take full advantage of PGPB inoculants.

---

**Supplementary Figure 1.**
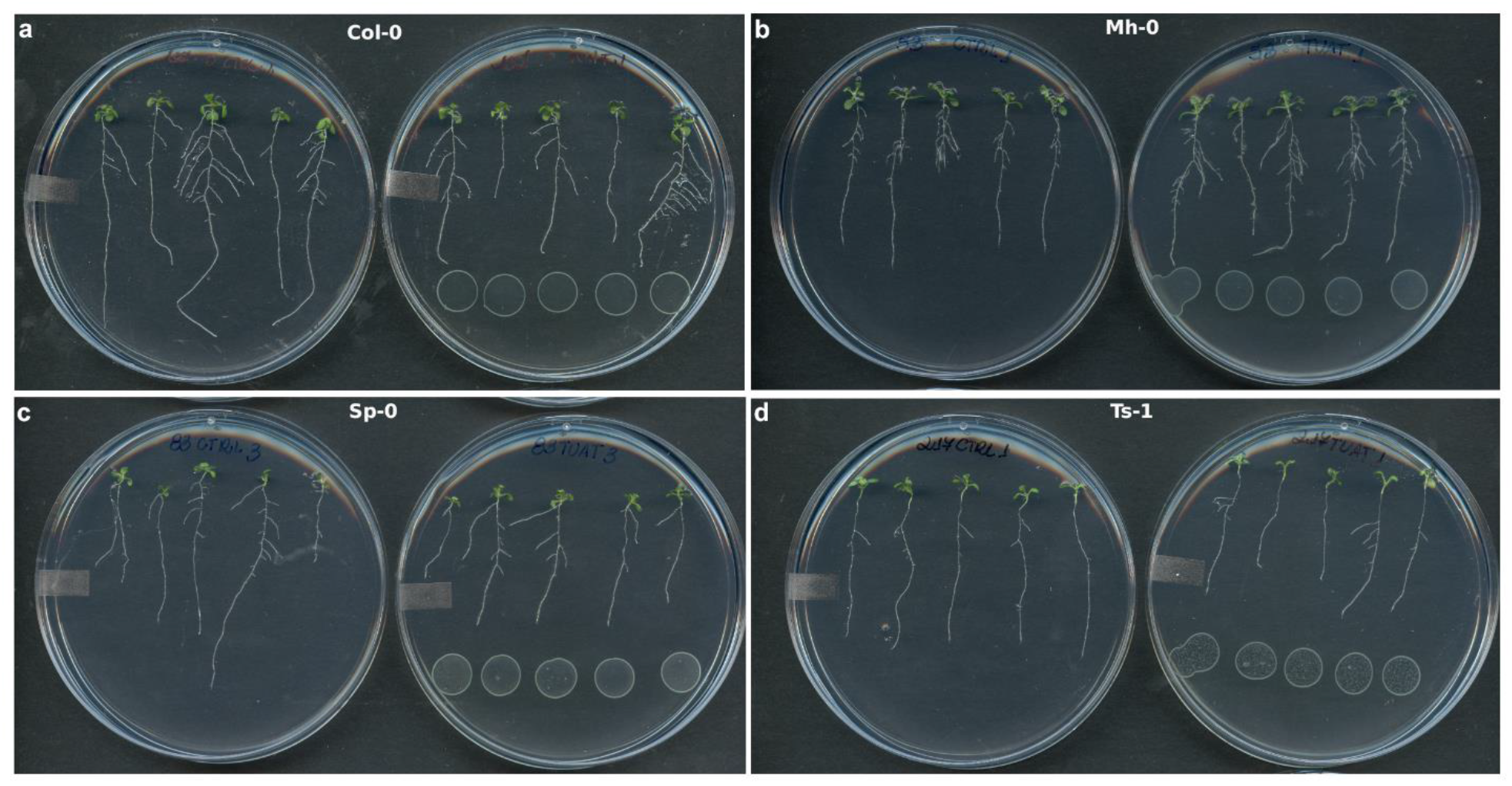
A. thaliana response upon TUAT-1 inoculation. (a) is the reference ecotype, Col-0. (b), (c) and (d) are positive (Mh-0), no (Sp-0) and negative (Ts-1) responsive ecotypes, respectively.

**Supplementary Figure 2.**
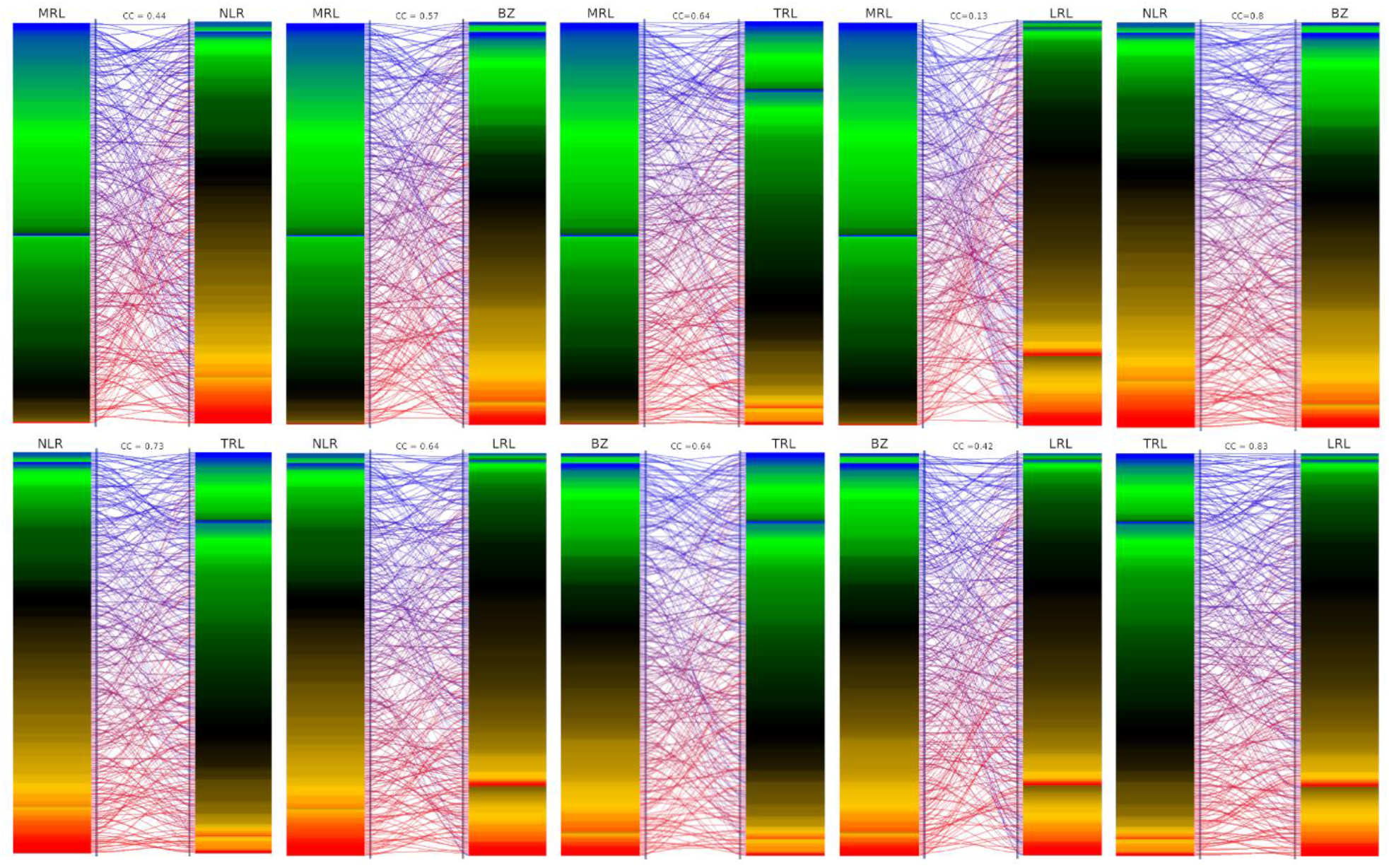
Correlation between traits Δ phenotypes values for each ecotype. The phenotype correlation coefficient (CC) value is above each pair of traits. The heatmaps were built sorting from the lowest phenotype Δ to the highest and then separated by the plant response category for each trait. Lines between the columns connects the same ecotype.

**Supplementary Figure 3.**
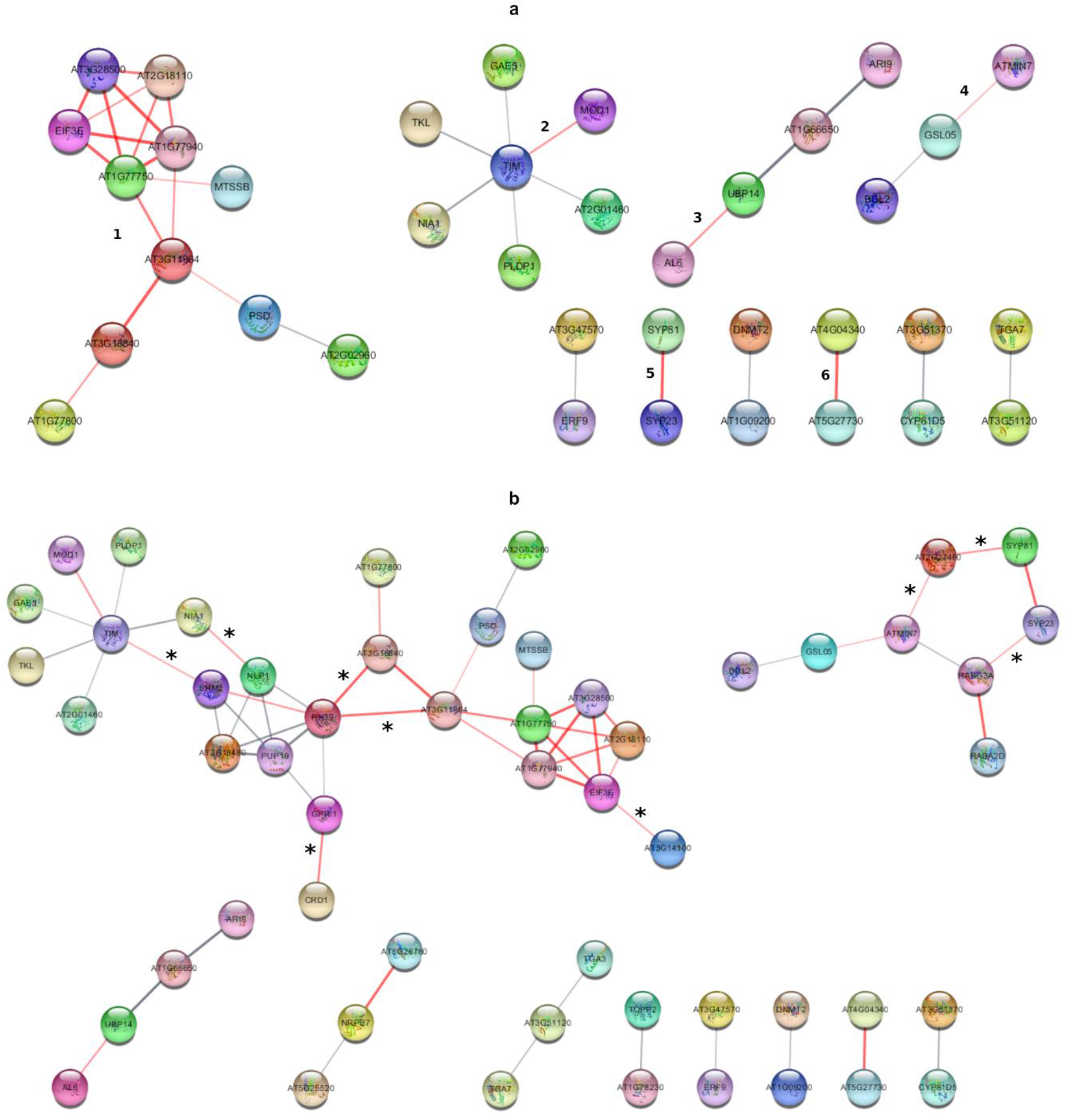
Gene interaction networks. (a) Gene interaction network built from all identified genes (AE and RE data set). Numbers close to the genes connected by red lines indicate them number of the co-expressed groups or pairs. (b) Gene interaction network from all genes identified in this work and genes previous reported for Pseudomonas simiae WCS417r and Arabidopsis thaliana interaction GWAS (WINTERMANS, 2016). The thicker the line connecting the nodes, the strongest is the relation between them. The types of interactions evidence are known interactions from curated databases or experimentally determined; predicted interactions of genes neighborhood, fusions or co-occurrence; or genes reported in the same study, co-expressed genes or protein homology. Red lines connecting genes indicate co-expression. *co-expression between identified genes for TUAT-1 and WCS417r and Arabidopsis thaliana interaction.

**Supplementary Figure 4.**
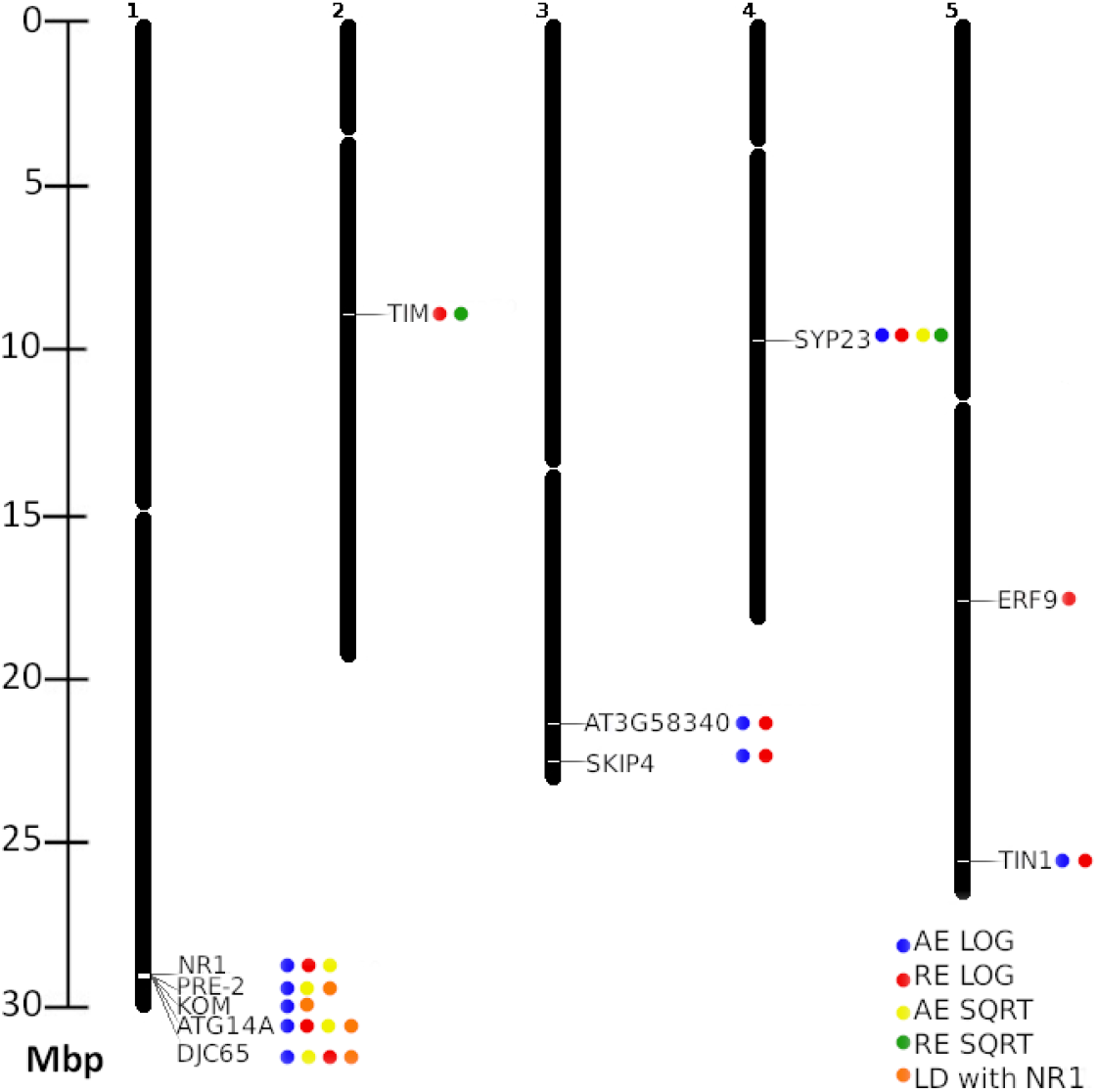
Chromosome map. Localization of the candidate genes in *A. thaliana* chromosomes. The circles next to the genes represent the analysis supporting the candidate gene selection.

**Supplementary Figure 5.**
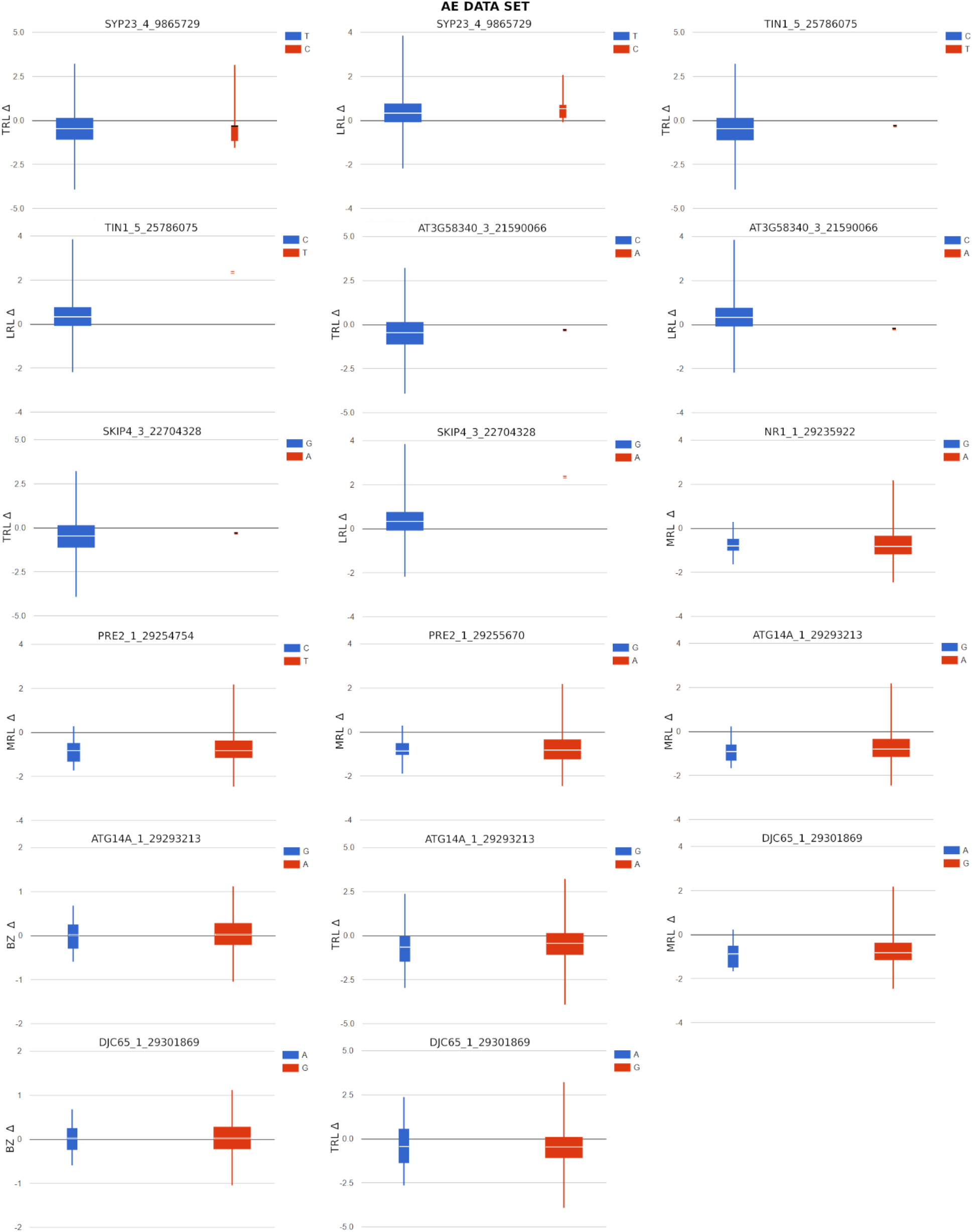
Allele effect of the candidate genes from the AE data set GWAS analysis.

**Supplementary Figure 6.**
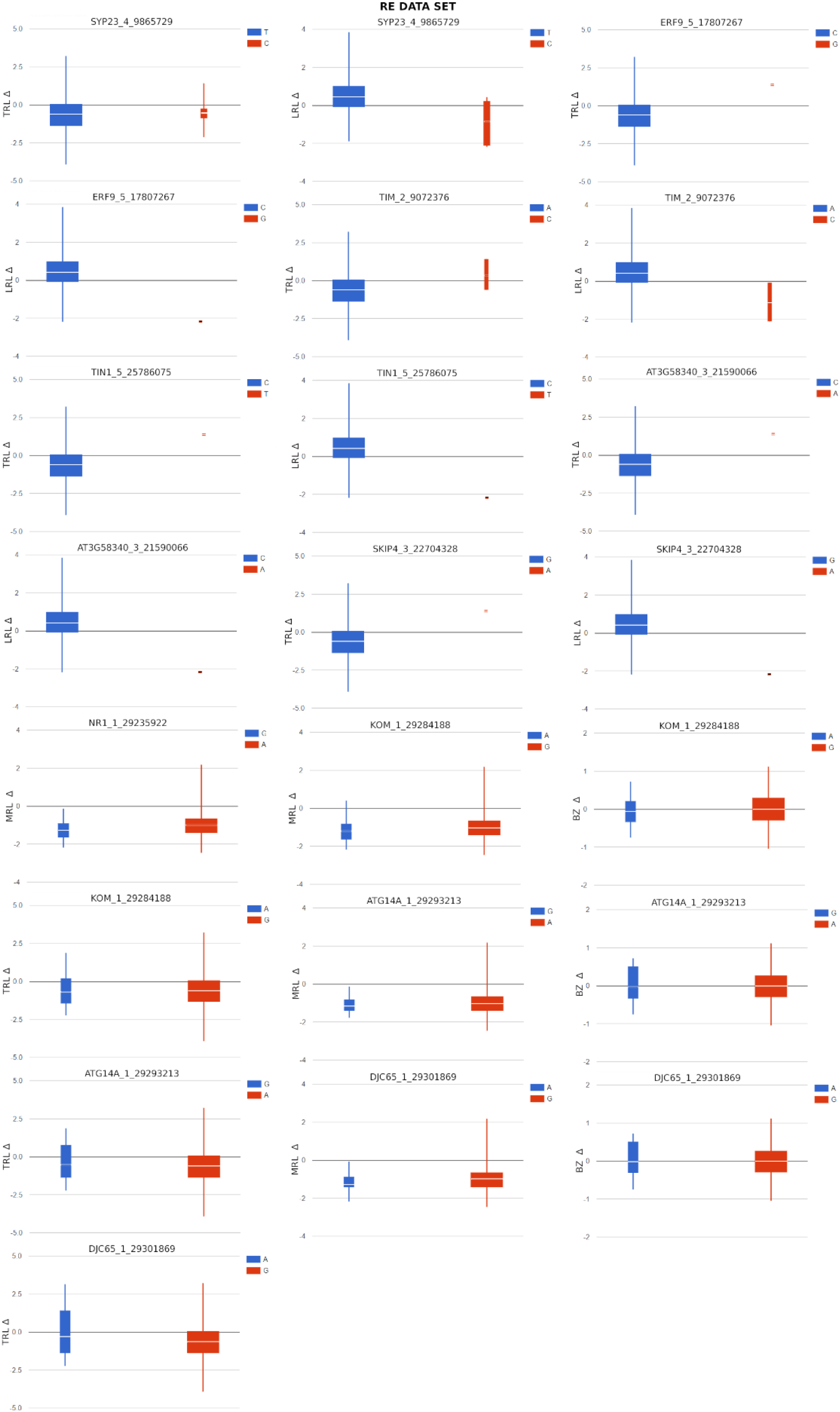
Allele effect of the candidate genes from the RE data set GWAS analysis.

## Notes

### Competing Interest Statement

The authors have declared no competing interest.

